# The influence of QTL allelic diversity on QTL detection in multi-parent populations: a simulation study in sugar beet

**DOI:** 10.1101/2020.02.04.930677

**Authors:** Vincent Garin, Valentin Wimmer, Dietrich Borchardt, Marcos Malosetti, Fred van Eeuwijk

## Abstract

Multi-parent populations (MPPs) are important resources for studying plant genetic architecture and detecting quantitative trait loci (QTLs). In MPPs, the QTL effects can show various levels of allelic diversity, which is an important factor influencing the detection of QTLs. In MPPs, the allelic effects can be more or less specific. They can depend on an ancestor, a parent or the combination of parents in a cross. In this paper, we evaluated the effect of QTL allelic diversity on the QTL detection power in MPPs.

We simulated: a) cross-specific QTLs; b) parental and ancestral QTLs; and c) bi-allelic QTLs. Inspired by a real application, we tested different MPP designs (diallel, chessboard, factorial, and NAM) derived from five or nine parents to explore the ability to sample genetic diversity and detect QTLs. Using a fixed total population size, the QTL detection power was larger in MPPs with fewer but larger crosses derived from a reduced number of parents. The use of a larger set of parents was useful to detect rare alleles with a large phenotypic effect. The benefit of using a larger set of parents was however conditioned on an increase of the total population size. We also determined empirical confidence intervals for QTL location to compare the resolution of different designs. For QTLs representing 6% of the phenotypic variation, using 1600 offspring individuals, we found 95% empirical confidence intervals of 50 and 26 cM for cross-specific and bi-allelic QTLs, respectively.

MPPs derived from less parents with few but large crosses generally increased the QTL detection power. Using a larger set of parents to cover a wider genetic diversity can be useful to detect QTLs with a reduced minor allele frequency when the QTL effect is large and when the total population size is increased.

## I. INTRODUCTION

The use of multi-parent populations (MPPs) for quantitative trait loci (QTLs) detection is growing in popularity. With respect to the bi-parental crosses, MPPs address a larger genetic diversity. With respect to the association panels, MPPs reduce the chance of false positive QTL detection due to a better control over the population structure [1]. Here, we focus on MPPs composed of bi-parental crosses without further intercrossing. This definition does not cover MPPs like the multi-parent advanced generation inter-cross (MAGIC) populations [2]. Different statistical procedures exist to detect QTLs in MPPs but generally those methods, like the one adapting models used in genome-wide association studies [3, 4], do not model properly the diversity of allelic effects present in MPPs. Similarly, most of the MPPs simulation studies have not simulated the wide range of QTL allelic effects present in those populations. Therefore, we investigated the QTL detection power in MPPs using scenarios that accounted better for the MPP QTL allelic diversity. We also determined empirical confidence intervals for the detected QTLs, which is an essential information for marker assisted selection (MAS) [5]. To the extent of our knowledge, no article provides such information in MPPs composed of crosses.

### MPP design

Many MPP designs have been evaluated through simulation studies [3, 6, 7, 8]. The nested association mapping (NAM) design is a collection of crosses between a central parent and peripheral lines [3] (Figure 1). In a diallel design, each *p* parent is crossed with the *p* − 1 other parents [9]. In a factorial design, a set of parents, A, is fully or partially crossed with another set of parents, B [9].

**Figure 1:**
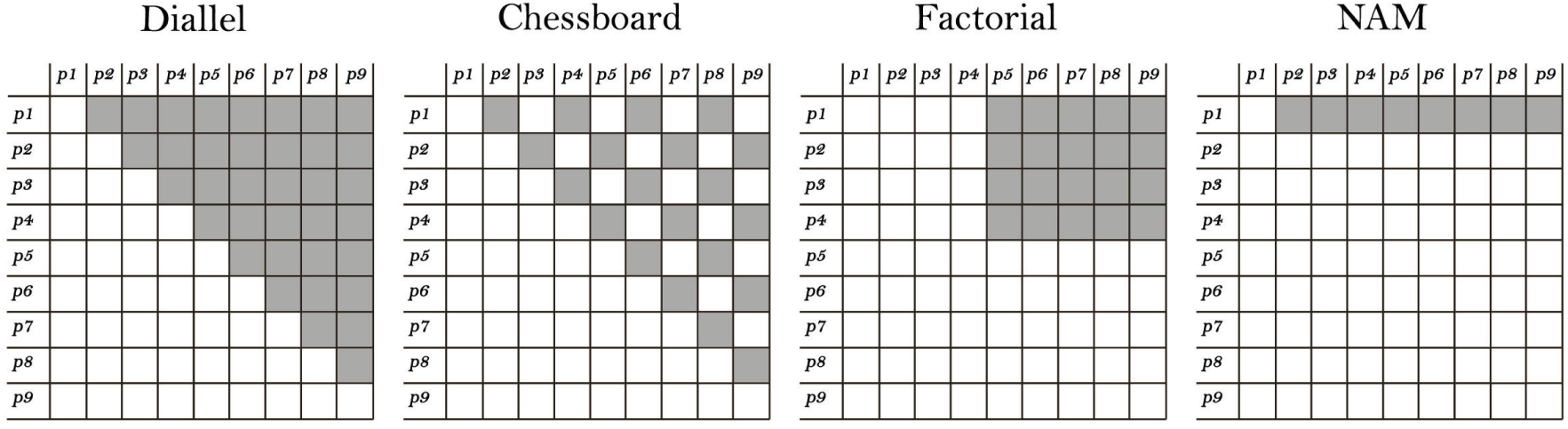
MPP designs illustration. Diallel: crosses between p and p − 1 parents. Chessboard: diallel structure without all possible crosses. Factorial: crosses between a set of parents A and another set of parents B. NAM: Crosses between a central parent and peripheral lines.

To define an MPP design, we can look at the crossing scheme, the number of parents or crosses, and the number of individuals per cross. A larger set of parents covers a wider genetic diversity by sampling more alleles. For a given amount of resources (fixed total population size), more parents implies more crosses and therefore reduces the number of individuals per cross. When more alleles segregate, each allele has a lower frequency in the population. Therefore, the number of parents represents a trade-off between the number of alleles sampled and the sample size to detect their effect.

According to the literature, MPP designs with a reduced number of large crosses are more powerful [10, 11, 6]. Some authors tried to determine an optimal number of parents and of individuals per cross. For example, in diallel and single round robin designs, [12] found that the optimal number of parents corresponded to crosses of 100 genotypes. In MPPs with a fixed population size, [8] determined analytically that the detection power was only influenced by the number of parents and not by the MPP design. She showed however that the power reached a plateau after six parents. The trade-off between the covered genetic diversity and the sample size to detect QTL allele is a question that deserves further investigation. The answer to this question could be influenced by the QTL allelic diversity present in MPPs.

### QTL allelic diversity

In MPPs, since the crosses are derived from multiple parents, more alleles can potentially segregate with effects that are more or less diverse/consistent. We define four types of QTL allelic effects from the most diverse and specific to the most consistent and shared. In an MPP, the QTL effects can be defined in terms of allele origin and/or mode of action (Figure 2). The first QTL allelic effect (cross-specific) represents an epistatic interaction between a parental allele and a cross genetic background [13]. Thus, cross-specific QTL effects can only be estimated within crosses (nested effects). The other QTL allelic effects are defined in terms of parental, ancestral, or single nucleotide polymorphism (SNP) alleles with consistent effects across crosses. The QTL allelic effects can be common because the alleles are unique to: 1) a common parent (parental), 2) a common ancestral line (ancestral), or 3) a common causal SNP (bi-allelic).

**Figure 2:**
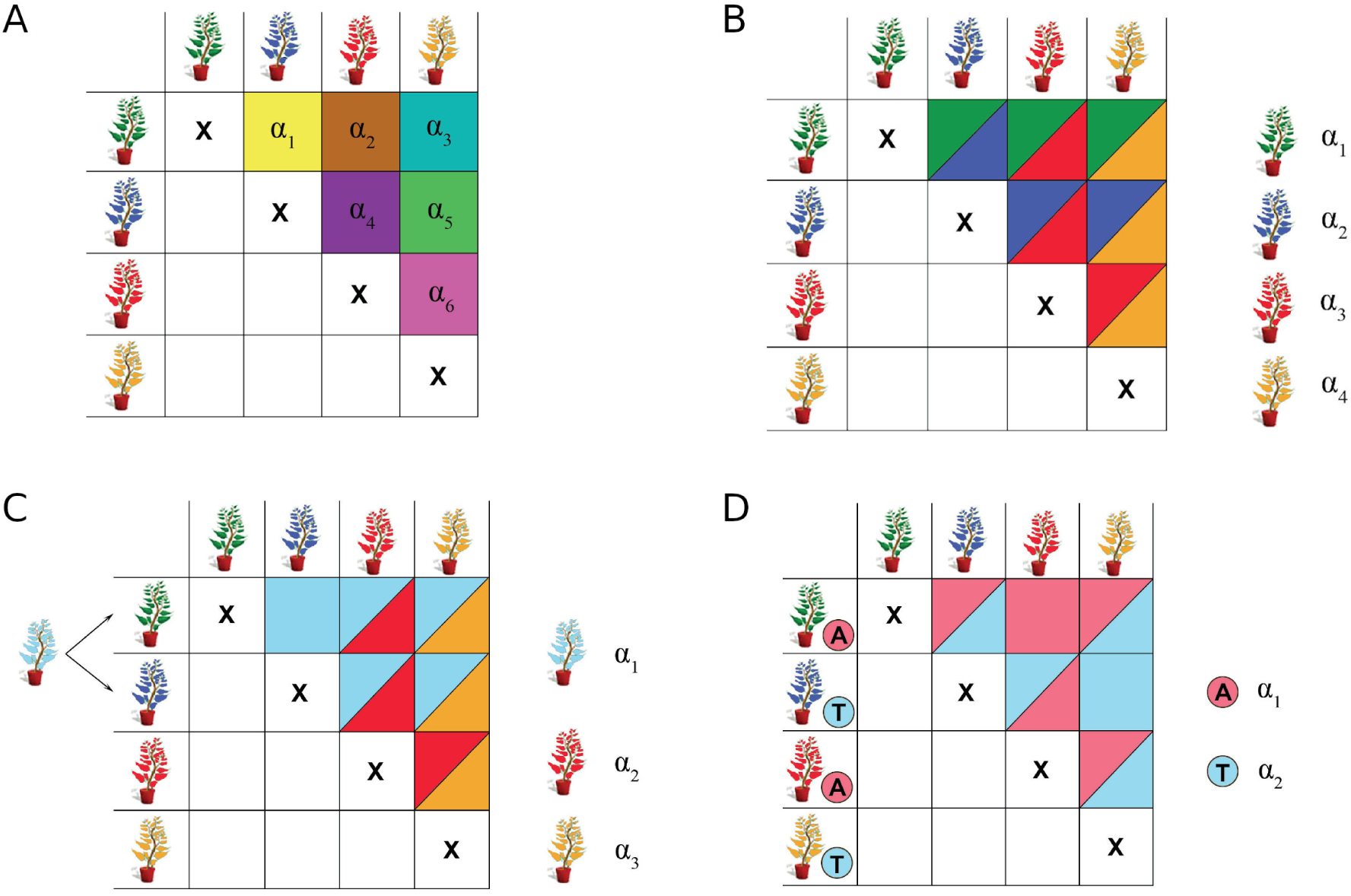
QTL effects illustration. A) cross-specific QTL: the allelic effects are different in each cross due to some interaction between a parental allele and the cross genetic background B) parental QTL: each parent carry a unique allele with consistent effect across the MPP C) ancestral QTL: parents inherit a reduced number of allele due to shared ancestors. Each ancestral allele has a consistent effect across the MPP D) bi-allelic QTL: parents with the same SNP marker score are assumed to inherit the same allele.

In the rest of the paper, we will refer to these four types of QTL effects calling them: cross-specific, parental, ancestral, and bi-allelic. Generally, the number of allelic effects that needs to be estimated decreases from the cross-specific to the bi-allelic QTLs. Therefore, the sample size to estimate the individual allelic or cross-specific QTL effects increases from the cross-specific to the bi-allelic QTLs. We hypothesize that the QTL allelic diversity has a strong influence on the detection of QTL in MPPs.

### Statistical models

In MPPs, the choice of the statistical model used for QTL detection should account for the design properties and the variety of QTL effects. [11], [10] and [6] assumed cross-specific QTL effects, which statistically corresponds to a saturated model. [8] and [14] considered the connection between crosses due to common parents using consistent parental QTL effects. [7] and [15] estimated multi-allelic QTL effects with an allele number between two and the number of parents. [12] used bi-allelic QTL models similar to the ones used in association studies. The Bayesian approach proposed by [16] is an elegant solution that can estimate the number of alleles and the global QTL variance. However, it also suffers from limitations because, according to the authors, with 600 individuals, it can only distinguish five alleles effectively. The computation in a Bayesian approach can also be intensive.

To capture the MPP QTL effect diversity, we used models assuming the different cross-specific, parental, ancestral and bi-allelic QTLs defined previously. We ran simulations based on genetic models with different levels of QTL allelic diversity that could better represent the genetic architectures present in MPPs. We evaluated the performance of our models on four common MPP designs with five or nine parents to explore the ability to sample genetic diversity for QTL detection. Besides that, we also evaluated the effects of the population size, the QTL allelic diversity, the QTL effect size, and the detection model on the detection power, the resolution, and the false discovery rate. The results allowed us to provide guidelines to design MPPs in sugar beet. Those guidelines could be useful for other MPPs or crops.

## II. METHODS

### MPP design

We evaluated the detection of QTLs on four MPP designs composed of *F*_2_ crosses : diallel, chessboard, factorial, and NAM (Figure 1). Given a fixed total population size, these designs represent different strategies to sample the QTL allelic diversity. The diallel design maximizes the number of estimable allele genetic background interactions (each parent *p* is used in *p* − 1 crosses) but limits the number of individuals per cross. The chessboard design is a compromise that samples a reduced number of allele by background interactions (each *p* parent is used in *p*/2 or [(*p* − 1)/2] ± 1 crosses) but allows more individuals per cross. The factorial design can be useful to cross two contrasting sets of parents (e.g. donors and recipients). Finally, the NAM design can be used to explore allelic diversity with respect to a reference parent [17]. In our case, the NAM design had the largest cross size.

We simulated the MPP designs using the genotypic data from nine parents coming from the sugar beet breeding program of KWS SAAT SE. Six parents were almost fully inbred with < 1% of heterozygous markers and three were partially inbred with around 18% of heterozygous markers. The use of an existing set of parents provided a realistic basis to our simulations in terms of genetic properties. We simulated a reference diallel population composed of the 36 possible *F*_2_ crosses between the nine parents. Each cross contained 450 genotypes, with recombination and meiosis simulated using a random Poisson process based on the genetic map. For each *in silico* QTL mapping experiment, we simulated the QTL effects on the reference population and sampled the genotypes from that population to form realizations of the tested MPP designs. We fixed the total population size to 800 or 1600 and ‘crossed’ either five or nine parents. The number of crosses varied from 4 to 36 and the cross size from 22 to 400 individuals (Table 1).

**Table 1:**
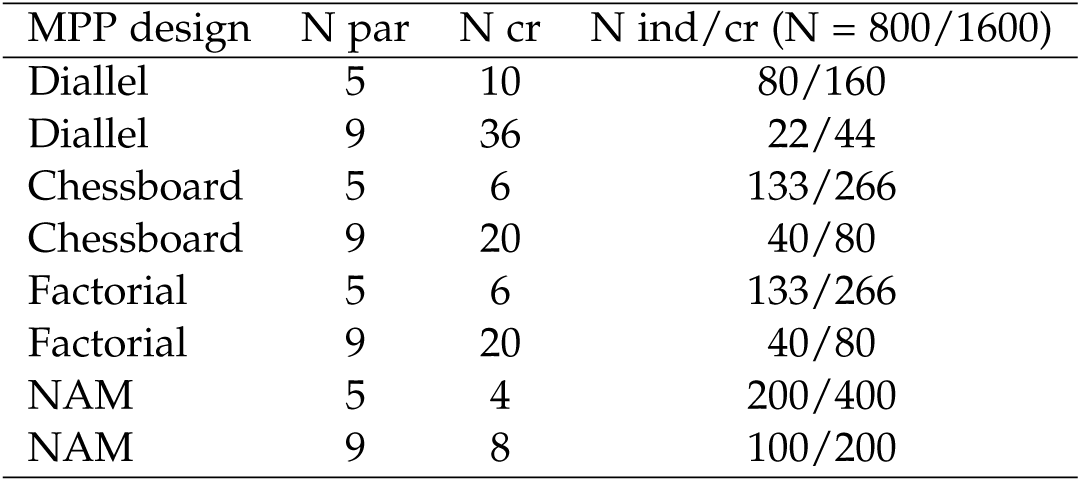
MPP designs properties with the number of parents (N par), the number of crosses (N cr) and the number of individuals per crosses (N ind/cr).

### QTL genetic model

The QTL effects were simulated from a genetic map containing 5000 SNP markers spread across nine chromosomes. The average minor allele frequency (MAF) was 0.29 with values between 0.04 and 0.5. The average genetic distance between markers was equal to 0.2 cM with a maximum of 10.6 cM. We simulated seven types of QTLs (Table 2 and Supplementary file S1). Q1 and Q2 were cross-specific and had non-zero allelic effects in half and one third of the crosses, respectively. Setting some cross-specific effects to zero implies that the QTL does not interact with those backgrounds, which seems a reasonable assumption to make from our experience of analyzing real data. Q3 and Q4 were parental. Q3 had a different allelic effect for each parents, while Q4 only had a single non-zero parental allelic effect randomly assigned. Q5 and Q6 were ancestral, with ancestry groups determined by clustering the nine parental lines on local genetic similarity in a 10 cM window using the R package clusthaplo [14]. On average, we detected 3.9 ancestral alleles along the genome. While Q5 had a different allelic effect for each ancestral group, Q6 only had a single non-zero ancestral allelic effect randomly assigned. Q4 and Q6 were bi-allelic QTLs with a parental and ancestral basis. The last QTLs (Q7) were bi-allelic with an effect attached to the minor SNP allele.

**Table 2:** Simulated QTLs effects described by type of allelic effect, segregation, number of alleles or QTL effects, and QTL genetic model.

| QTL | Allelic eff. | Segregation | N all. (Q. eff.) | Gen. mod. |
| --- | --- | --- | --- | --- |
| Q1 | cr. sp. | 1/2 of the cr. | 18 | M1 (M5) |
| Q2 | cr. sp. | 1/3 of the cr. | 12 | M1 (M5) |
| Q3 | parental | all parents | 9 | M2 (M5) |
| Q4 | parental | 1 parent | 2 | M2 (M5) |
| Q5 | ancestral | all ancestors | 4 | M3 (M5) |
| Q6 | ancestral | 1 ancestor | 2 | M3 (M5) |
| Q7 | bi-allelic | minor SNP allele | 2 | M4 (M5) |

We sampled the non-zero QTL allelic effects from a uniform distribution (min=1; max=10) with the signs randomly assigned with an equal probability. In each *in silico* QTL mapping experiment, we simulated genetic models with eight QTLs located on different chromosomes keeping one chromosome free to control the false discovery rate. We simulated four QTLs with a small effect and four with a big effect representing 2% and 6% of the phenotypic variation, respectively. The total genetic contribution was always equal to 32% of the phenotypic variance. The sampled QTL allelic values were standardized to have realized QTL variances equal to the defined ones. The remaining phenotypic variance representing the environmental and plot error was simulated using a normal distribution. For details about the phenotype simulation see Supplementary file S2.

We simulated five QTL genetic models (M1-M5). The first four models used only QTLs with a single type of allelic effect. M1 contained only cross-specific QTLs (Q1 and Q2), M2 only parental QTLs (Q3 and Q4), M3 only ancestral QTLs (Q5 and Q6), and M4 only bi-allelic QTLs (Q7). In the last multi-QTL effects (MQE) genetic model (M5), a combination of all QTL effects was used. It contained Q1 to Q6 and two times Q7 to have two QTLs of each type. Those genetic models, especially the MQE model, should be more representative of the QTL effects diversity present in MPPs than the one generally used in MPP simulations. In the rest of the paper, we will refer to the QTL genetic models calling them cross-specific, parental, ancestral, bi-allelic and MQE genetic models or we will refer directly to their types of QTL allelic effects.

### QTL detection model and procedure

The QTL detection models had the following form:

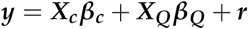

where ***X***_***c***_ represented a cross-specific intercept. ***X***_***Q***_ and ***β***_***Q***_ represented the QTL incidence matrix and the QTL effects that varied according to the number of QTL alleles or estimated effects [18]. We evaluated the QTL detection performances of four models assuming: cross-specific, parental, ancestral and bi-allelic QTL allelic effects. The residual term ***r*** followed the linear model assumption of normality with a homogeneous variance 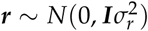.

The QTL detection procedure was composed of a simple interval mapping scan to select cofactors. On each chromosome, we selected positions with − *log*10(*p* − *value*) >= 4. We applied an exclusion window of 100 cM around the selected cofactors and iterated until no remaining position could be chosen. The large exclusion window allowed us to reduce the number of cofactors per chromosome to avoid model overfitting. Then, using the cofactors we performed a composite interval mapping to build a multi-QTL model. The QTL positions were selected with a minimum distance of 30 cM using the same procedure and threshold used for cofactors selection. The final list of QTLs was evaluated using a backward elimination.

The choice of a − *log*10(*p* − *value*) >= 4 threshold was based on values determined by permutation in real data analyses. For example, in [19], the threshold values varied between 3.4 to 5.6 for a type I error of 10%. The QTL detection scans were performed using the R package mppR [20].

### Evaluation statistics

We evaluated the QTL detection power calculating the true positive rate (TPR) as the fraction of correctly detected QTLs assuming a maximum distance of 5, 10 and 20 cM between the simulated QTL and the detected position. The TPR at the whole chromosome level (TPR chr) was the fraction of detected QTL without minimum distance with respect to the simulated QTL position. We calculated the corresponding false discovery rate (FDR) as the proportion of detected QTLs that were distant from a simulated QTL position by more than 5, 10 and 20 cM. The false discovery rate on the chromosome without simulated QTL (FDR chr) was the percentage of runs where a QTL was wrongly detected. The TPR and the FDR did not sum to one because they were calculated with different denominators. The TPR denominator was the number of simulated QTLs (8) and the FDR denominator was the number of detected QTLs.

We evaluated the resolution of the QTL detection (dQTL) by measuring the distance between a simulated QTL and the largest significant peak on the chromosome. This information allowed us to determine a distribution for dQTL and to provide confidence intervals (CI) for QTL positions detected in MPPs composed of *F*_2_ crosses. We defined the empirical CI as *CI*_*α*_ = 2 ∗ *dQTL*_*α*_, where *dQTL*_*α*_ is the *α* quantile of the dQTL distribution. The value *dQTL*_*α*_ is multiplied by two to account for situations where the detected QTL is positioned on the left or the right side of the true QTL. *CI*_*α*_ is the interval around the QTL position that contains the true QTL position with an *α* probability.

To evaluate the effect of the different parameters, we computed analyses of variance (ANOVAs) with the TPR at 10 cM as response and as explanatory factors: the MPP design (D), the number of parents (N), the QTL detection model (M), the QTL size (Qs), and the QTL effect (Qe) (Model 1). We included in the model the two-way interactions between the MPP design, the number of parents, the QTL detection model and the QTL size. The error term was normally distributed 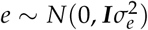.

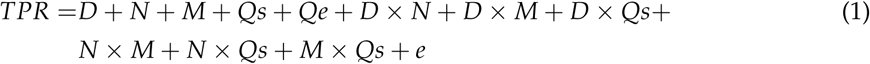

We interpreted the ANOVA results by calculating least-squares means (LSMs), following a model selection procedure that retained significant interactions and significant main effects as well as non-significant main effects that underlie significant interactions. LSMs allowed us to get predictions averaged over a set of factors to understand the effects of those factors on the TPR. Model 1 evaluates jointly the effect of all parameters and determine their relative contribution while, in most of the MPP simulations, the influence of each parameter was analyzed one by one. We computed the LSMs using the R package emmeans [21].

We performed 50 replications of each *in silico* QTL mapping experiment for a different QTL genetic model (M1-M5). For each replication, we sampled MPPs given the population sizes (N = 800 or N = 1600), the MPP design (diallel, chessboard, factorial and NAM), and the number of parents (five or nine). On each sampled MPP, we performed QTL detection by the cross-specific, parental, ancestral and bi-allelic models. It represented a total of 16,000 calculated QTL profiles with 128,000 simulated QTL positions.

## III. RESULTS

### Global measurements

Table 3 contains the average TPR and FDR at 5, 10, and 20 cM, and the average dQTL per population size and per QTL genetic model. The results are averaged over all other factors (MPP design, number of parents, QTL detection model, QTL size, and QTL type). The TPR increased when the tolerance distance to the simulated QTL increased. However, the increase between the TPR at 20 cM and TPR chr (no minimum distance) was limited.

**Table 3:** Average TPR FDR and dQTL results. The TPR and FDR are measured at 5, 10 and 20 cM. TPR chr is the TPR with no maximal distance to the true QTL. FDR chr is the average FDR on the chromosome with non simulated QTL. The results are presented per population size (N = 800 and N = 1600) and QTL genetic model (cross-specific, parental, ancestral, bi-allelic, and MQE).

| Genetic model | N = 800 |  |  |  |  | N = 1600 |  |  |  |  |
| --- | --- | --- | --- | --- | --- | --- | --- | --- | --- | --- |
|  | Cr. sp. | Par. | Anc. | Biall. | MQE | Cr. sp. | Par. | Anc. | Biall. | MQE |
| TPR (%) (5cM) | 24 | 16 | 25 | 26 | 23 | 42 | 34 | 48 | 50 | 42 |
| FDR (%) (5cM) | 45 | 47 | 35 | 32 | 40 | 42 | 41 | 26 | 24 | 33 |
| TPR (%) (10cM) | 32 | 22 | 30 | 31 | 29 | 52 | 43 | 56 | 56 | 51 |
| FDR (%) (10cM) | 28 | 28 | 21 | 19 | 24 | 28 | 24 | 14 | 13 | 20 |
| TPR (%) (20cM) | 38 | 27 | 35 | 35 | 34 | 61 | 51 | 61 | 61 | 58 |
| FDR (%) (20cM) | 12 | 11 | 8 | 7 | 10 | 13 | 10 | 6 | 5 | 9 |
| TPR chr (%) | 41 | 30 | 37 | 37 | 37 | 66 | 56 | 64 | 63 | 62 |
| FDR chr (%) | 0.3 | 0.9 | 0.2 | 0.6 | 0.5 | 1.7 | 0.9 | 0.1 | 0.6 | 0.7 |
| dQTL (cM) | 7.1 | 8 | 6.1 | 5.3 | 7.1 | 6.5 | 6.8 | 4.3 | 3.9 | 5.4 |

FDR chr varied between 0.1 and 1.7%. On the chromosomes where QTLs were simulated, the FDRs were larger (e.g. at 10 cM around 25% and 20% for the N = 800 and N = 1600 populations respectively). The FDR decreased from the cross-specific and parental QTLs to the ancestral and bi-allelic ones. This tendency was more pronounced in the N = 1600 populations. For example, at 10 cM, the FDR was equal to 28%, 24%, 14% and 13% for the cross-specific, parental, ancestral, and bi-allelic genetic models, respectively.

dQTL followed a trend similar to the FDR and decreased from the cross-specific and parental QTLs to the ancestral and bi-allelic QTLs. For example, in the N = 1600 populations, dQTL decreased from 6.5 cM to 3.9 cM for the cross-specific and bi-allelic QTLs, respectively. In Table 4, we calculated the empirical CI for the 0.9, 0.95, and 0.99 quantile values of the dQTL distribution per QTL and population size for each genetic model. For an illustration of the dQTL distributions, see Supplementary file S3. Table 4 gives an estimation of the QTL CI. For example, in a population of N = 800, with a mix of QTL effects (MQE) explaining each 6% (2%) of the phenotypic variation, the 95% empirical CI was equal to 46 cM (70 cM). Finally, we noticed that increasing the population size from 800 to 1600 leaded to a smaller reduction of dQTL for the cross-specific and the parental QTLs than for the ancestral and bi-allelic ones. In general, the TPR, FDR, and dQTL results obtained for the MQE genetic model seemed to be an average of the individual types of genetic model.

**Table 4:**
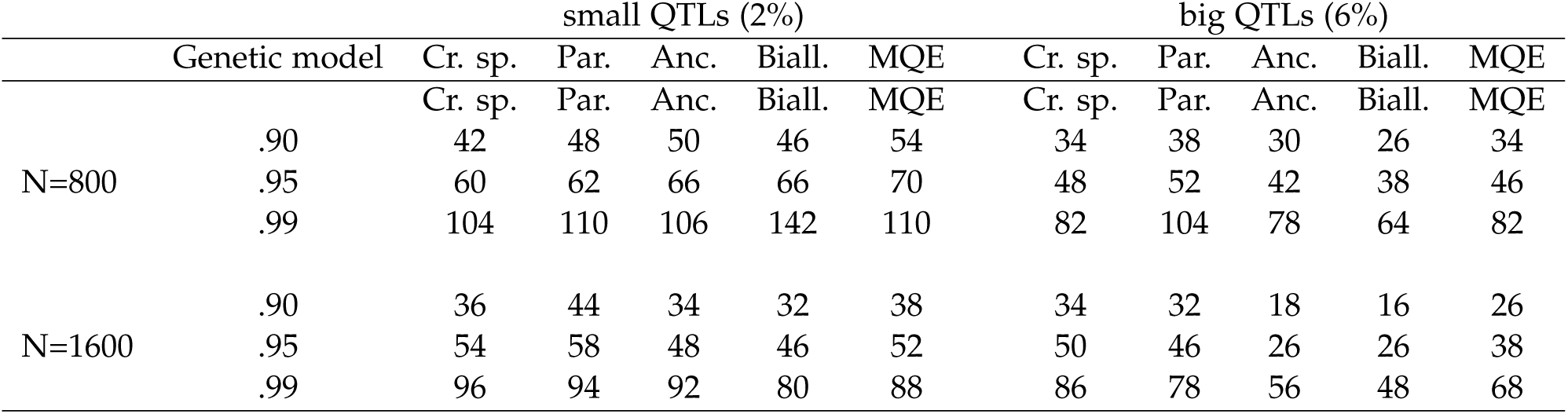
Empirical confidence interval in cM around the detected position based on the 90, 95 and 99 % quantile value from the dQTL distribution per QTL size (small 2% and big 6%), per QTL genetic model (cross-specific, parental, ancestral, bi-allelic, and MQE), and per population size (N = 800 and N = 1600).

|  |  | small QTLs (2%) |  |  |  |  | big QTLs (6%) |  |  |  |  |
| --- | --- | --- | --- | --- | --- | --- | --- | --- | --- | --- | --- |
|  | Genetic model | Cr. sp. | Par. | Anc. | Biall. | MQE | Cr. sp. | Par. | Anc. | Biall. | MQE |
| N=800 | .90 | 42 | 48 | 50 | 46 | 54 | 34 | 38 | 30 | 26 | 34 |
|  | .95 | 60 | 62 | 66 | 66 | 70 | 48 | 52 | 42 | 38 | 46 |
|  | .99 | 104 | 110 | 106 | 142 | 110 | 82 | 104 | 78 | 64 | 82 |
| N=1600 | .90 | 36 | 44 | 34 | 32 | 38 | 34 | 32 | 18 | 16 | 26 |
|  | .95 | 54 | 58 | 48 | 46 | 52 | 50 | 46 | 26 | 26 | 38 |
|  | .99 | 96 | 94 | 92 | 80 | 88 | 86 | 78 | 56 | 48 | 68 |

### MPP design

Tables 5 and 6 contain the ANOVA F-statistics describing the effect of the MPP design, the number of parents, the QTL detection model, the QTL size, and the QTL effect on the TPR at 10 cM for the N = 800 and N = 1600 populations, respectively. Those results show that the MPP design was mostly influential for the cross-specific QTLs. The MPP design F-statistic of the cross-specific genetic model was equal to 91.9 and 44.8 in the N = 800, and N = 1600 populations, respectively. For the other QTL genetic models, except the MQE, the maximum MPP design F-statistic was equal to 4.4. In Figure 3, we plotted the LSM of TPR over the MPP design per QTL size. We could again observe that the cross-specific QTLs were the only QTL effects for which the MPP design influences the TPR. For the cross-specific QTLs, the TPR increased from the diallel to the NAM design.

**Table 5:** ANOVA degrees of freedom (df), and F-statistics with significance of the MPP design, the number of parents, the QTL detection model, the QTL size and the QTL effect on TPR per QTL genetic model (cross-specific, parental, ancestral, bi-allelic, and MQE) for the N = 800 populations. Adjusted and cross-validation R2 of the models.

|  | df | Cr. sp. |  | Parental |  | Ancestral |  | Bi-allelic |  | MQE |  |
| --- | --- | --- | --- | --- | --- | --- | --- | --- | --- | --- | --- |
| MPP des | 3 | 91.9 | *** | 2.6 |  | 1.4 |  | 3.4 | * | 15.6 | *** |
| Nb. par | 1 | 188.6 | *** | 131.8 | *** | 37 | *** | 22.4 | *** | 97 | *** |
| Det. model | 3 | 75.5 | *** | 8.9 | *** | 87.4 | *** | 99 | *** | 5 | ** |
| QTL size | 1 | 693.3 | *** | 845.1 | *** | 1372.9 | *** | 1167.8 | *** | 824 | *** |
| QTL type | 0 – 3<sup>a</sup> | 8.5 | ** | 0 |  | 0 |  |  |  | 10.9 | *** |
| MPP des x Nb. par | 3 | 9.7 | *** | 1.8 |  | 1.3 |  | 0.1 |  | 3.4 | * |
| MPP des x Det. model | 9 | 4.5 | *** | 1.2 |  | 1.9 |  | 1.5 |  | 0.5 |  |
| MPP des x QTL size | 3 | 2.4 |  | 3.1 | * | 11.2 | *** | 0.8 |  | 0.3 |  |
| Nb. par x Det. model | 3 | 3.2 | * | 2.5 |  | 13.9 | *** | 13.3 | *** | 2.3 |  |
| Nb. par x QTL size | 1 | 5.3 | * | 22 | *** | 1.1 |  | 1.4 |  | 3.5 |  |
| Det. model x QTL size | 3 | 22 | *** | 6.3 | *** | 17.8 | *** | 17.3 | *** | 0.2 |  |
| Residuals | 94 – 97<sup>a</sup> |  |  |  |  |  |  |  |  |  |  |
| adj. R2 (%) |  | 92 |  | 89 |  | 93 |  | 96 |  | 67 |  |
| CV R2 (%) |  | 90 |  | 85 |  | 91 |  | 92 |  | 65 |  |

\* p < 0.05; \*\* p < 0.01; \*\*\* p < 0.001
a. The df of the QTL type term is equal to 1, 1, 1, 0, 3 from the Cr. sp. to the MQE genetic model. The df of the residuals change according to the df of the QTL type term.

**Table 6:**
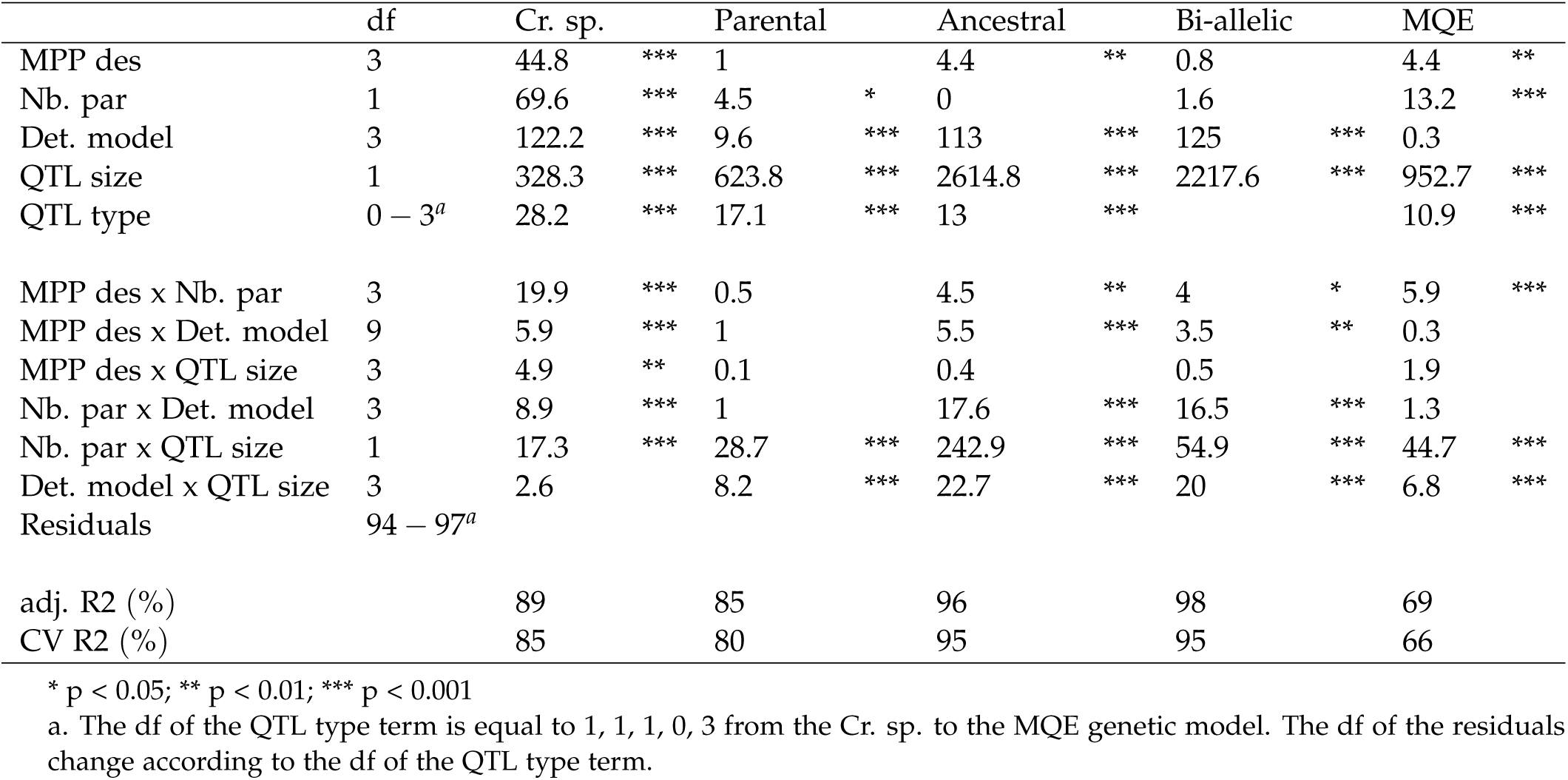
ANOVA degrees of freedom (df), and F-statistics with significance of the MPP design, the number of parents, the QTL detection model, the QTL size and the QTL effect on TPR per QTL genetic model (cross-specific, parental, ancestral, bi-allelic, and MQE) for the N = 1600 populations. Adjusted and cross-validation R2 of the models.

|  | df | Cr. sp. |  | Parental |  | Ancestral |  | Bi-allelic |  | MQE |  |
| --- | --- | --- | --- | --- | --- | --- | --- | --- | --- | --- | --- |
| MPP des | 3 | 44.8 | *** | 1 |  | 4.4 | ** | 0.8 |  | 4.4 | ** |
| Nb. par | 1 | 69.6 | *** | 4.5 | * | 0 |  | 1.6 |  | 13.2 | *** |
| Det. model | 3 | 122.2 | *** | 9.6 | *** | 113 | *** | 125 | *** | 0.3 |  |
| QTL size | 1 | 328.3 | *** | 623.8 | *** | 2614.8 | *** | 2217.6 | *** | 952.7 | *** |
| QTL type | 0 – 3<sup>a</sup> | 28.2 | *** | 17.1 | *** | 13 | *** |  |  | 10.9 | *** |
| MPP des x Nb. par | 3 | 19.9 | *** | 0.5 |  | 4.5 | ** | 4 | * | 5.9 | *** |
| MPP des x Det. model | 9 | 5.9 | *** | 1 |  | 5.5 | *** | 3.5 | ** | 0.3 |  |
| MPP des x QTL size | 3 | 4.9 | ** | 0.1 |  | 0.4 |  | 0.5 |  | 1.9 |  |
| Nb. par x Det. model | 3 | 8.9 | *** | 1 |  | 17.6 | *** | 16.5 | *** | 1.3 |  |
| Nb. par x QTL size | 1 | 17.3 | *** | 28.7 | *** | 242.9 | *** | 54.9 | *** | 44.7 | *** |
| Det. model x QTL size | 3 | 2.6 |  | 8.2 | *** | 22.7 | *** | 20 | *** | 6.8 | *** |
| Residuals | 94 – 97<sup>a</sup> |  |  |  |  |  |  |  |  |  |  |
| adj. $R^2$ (%) | | 89 | | 85 | | 96 | | 98 | | 69 | |
| CV $R^2$ (%) | | 85 | | 80 | | 95 | | 95 | | 66 | |

\* $p < 0.05$ ; \*\* $p < 0.01$ ; \*\*\* $p < 0.001$
a. The df of the QTL type term is equal to 1, 1, 1, 0, 3 from the Cr. sp. to the MQE genetic model. The df of the residuals change according to the df of the QTL type term.

**Figure 3:**
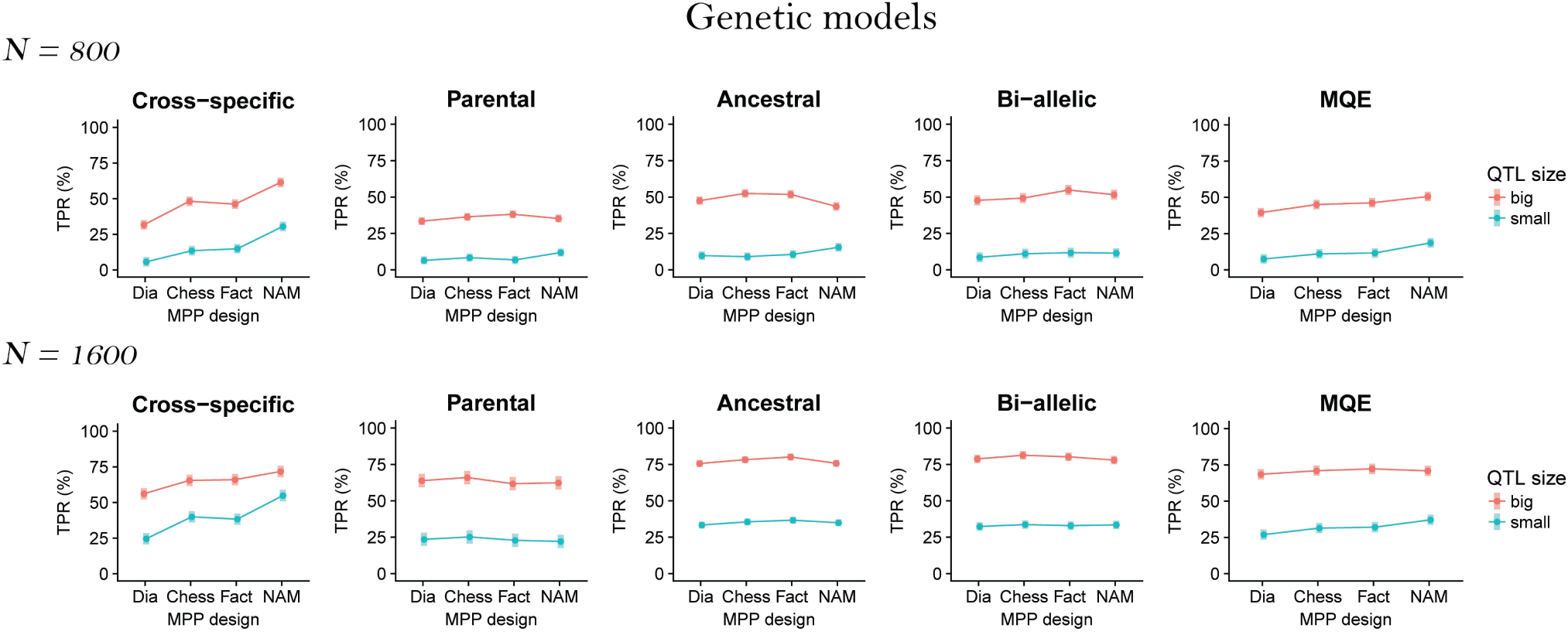
TPR LSM over the MPP designs (diallel, chessboard, factorial and NAM) and the QTL sizes (small 2% and big 6%) for all QTL genetic models (cross-specific, parental, ancestral, bi-allelic, and MQE) per population size (N = 800 and N = 1600).

### Number of parents

Concerning the number of parents used, we noticed in ANOVA Tables 5 and 6 that the number of parents main effect decreased from the cross-specific to the bi-allelic genetic model. For example, in the N = 800 populations, the F-statistic decreased from 188.6 to 22.4. In the N = 1600 populations, the number of parents main effect became non-significant for the ancestral and bi-allelic QTLs. In Figure 4, we plotted the TPR LSMs against the number of parents and the QTL size. We observed that generally the MPP designs with nine parents had a lower TPR. This trend was consistent in the N = 800 populations. However, in the N = 1600 populations, for the big QTLs, the TPR increased from five to nine parents for the parental, ancestral, and bi-allelic QTLs.

**Figure 4:**
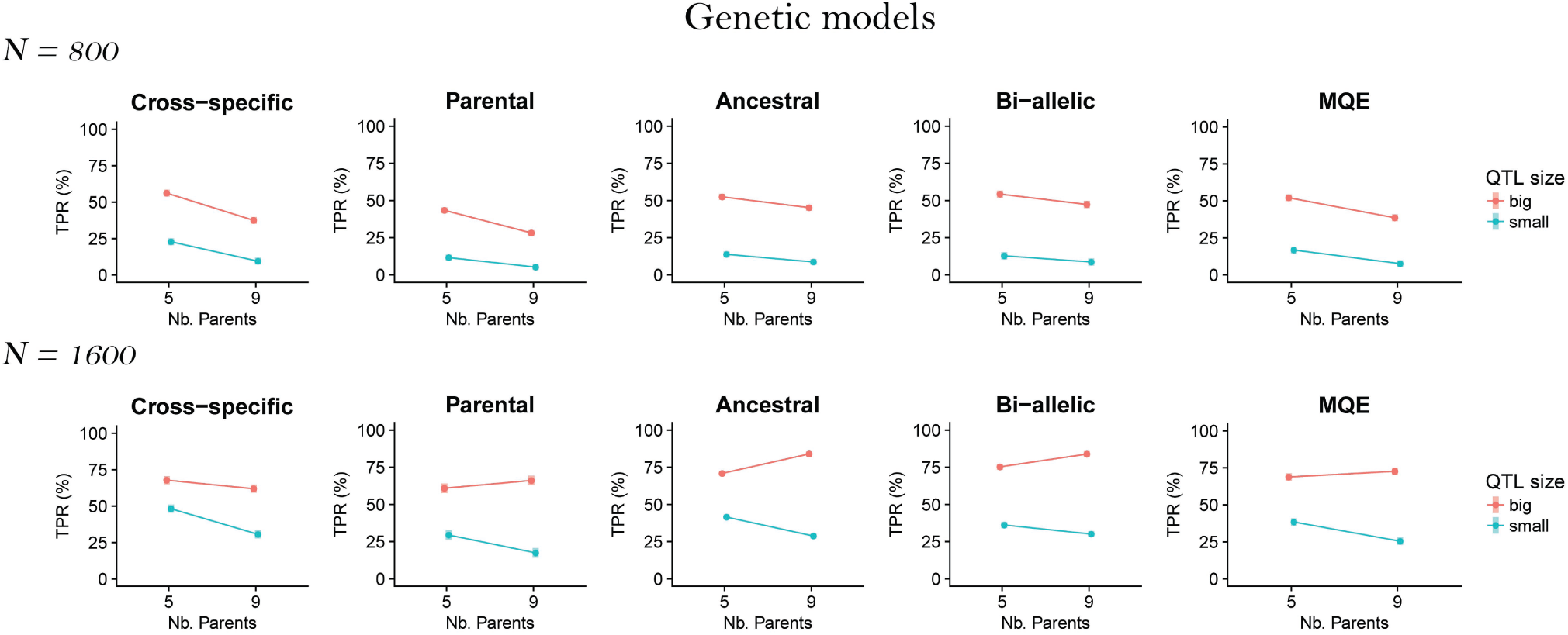
TPR LSM over the number of parents (5 and 9) and the QTL size (small 2% and big 6%) for all QTL genetic models (cross-specific, parental, ancestral, bi-allelic, and MQE) per population size (N = 800 and N = 1600).

We investigated in more details the situations where sampling nine parents increased the TPR. In Figure 5, we plotted the TPR LSMs against the number of parents for the big (6%) simulated QTLs (Q1 to Q7) in the N = 1600 populations. For the bi-allelic QTLs (Q7), we split the results in a low and a large MAF category given that the QTL MAF was below or above the median. For the parental, ancestral and bi-allelic effects we noticed that sampling a larger number of parents was more useful for the QTLs with a reduced number of QTL allelic effects and a reduced MAF (Q4, Q6 and Q7 low MAF). The best example is the difference between Q3 and Q4. Q3 had 9 parental alleles different from zero when Q4 only had one non-zero parental allele. The TPR of Q3 decreased while the TPR of Q4 increased when we sampled nine parents in place of five.

**Figure 5:**
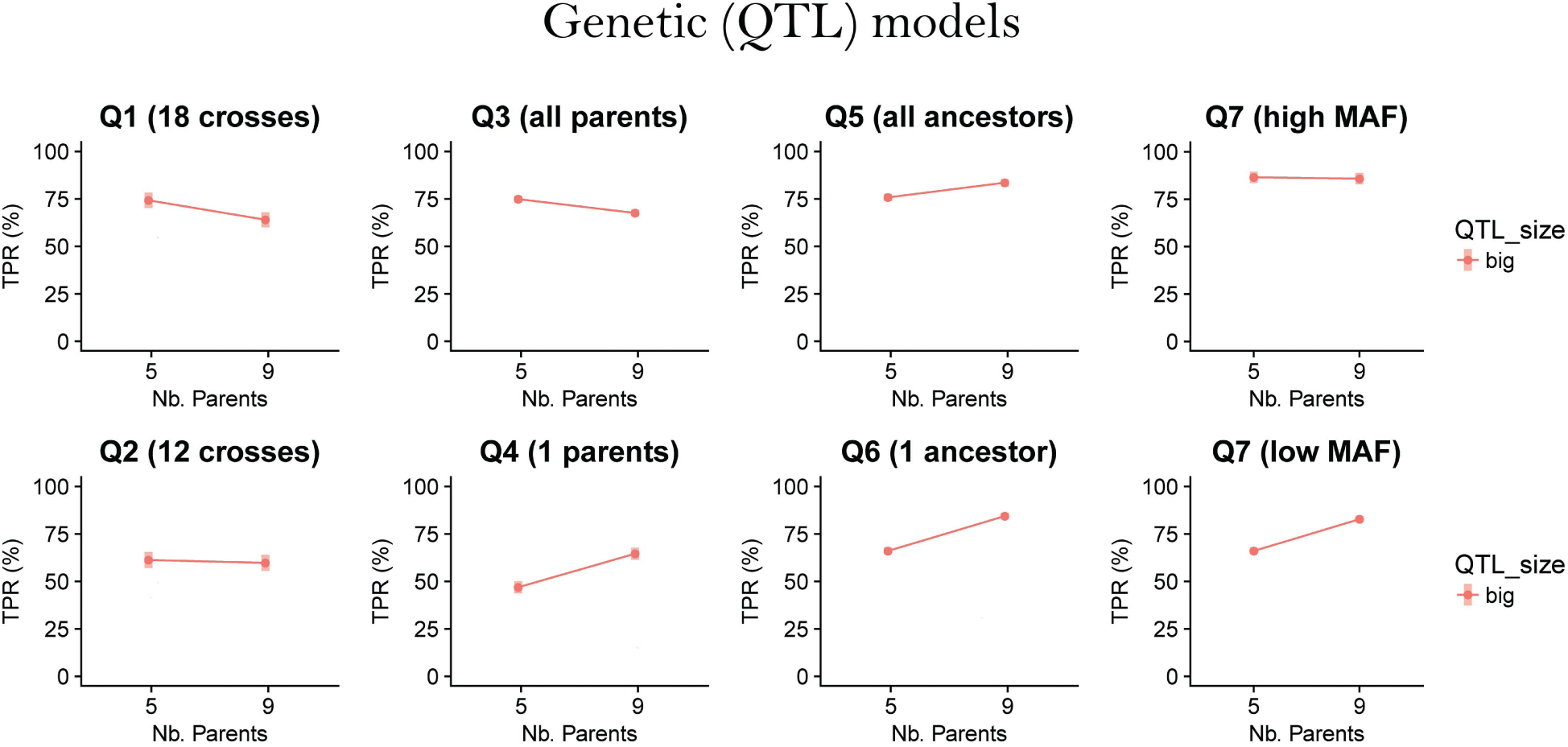
TPR LSM over the number of parents (5 and 9) for big (6%) simulated QTLs Q1-7 in populations N = 1600. The bi-allelic QTLs (Q7) were split into a low and a high MAF category given that their MAF was below or above the median.

### QTL detection model

The main effect for QTL detection model was highly significant for all QTL genetic models except for the MQE. For the MQE genetic model, the QTL detection model term was only moderately significant in the N = 800 populations. In Figure 6, we plotted the TPR LSMs against the QTL detection model and the QTL sizes. The QTL detection model effect was consistent with the way we simulated the QTLs. The QTL detection model that assumed the corresponding simulated QTL genetic model showed the best performance. For example, ancestral QTLs were detected with the largest TPR using an ancestral model. This result was stronger for the big QTLs.

**Figure 6:**
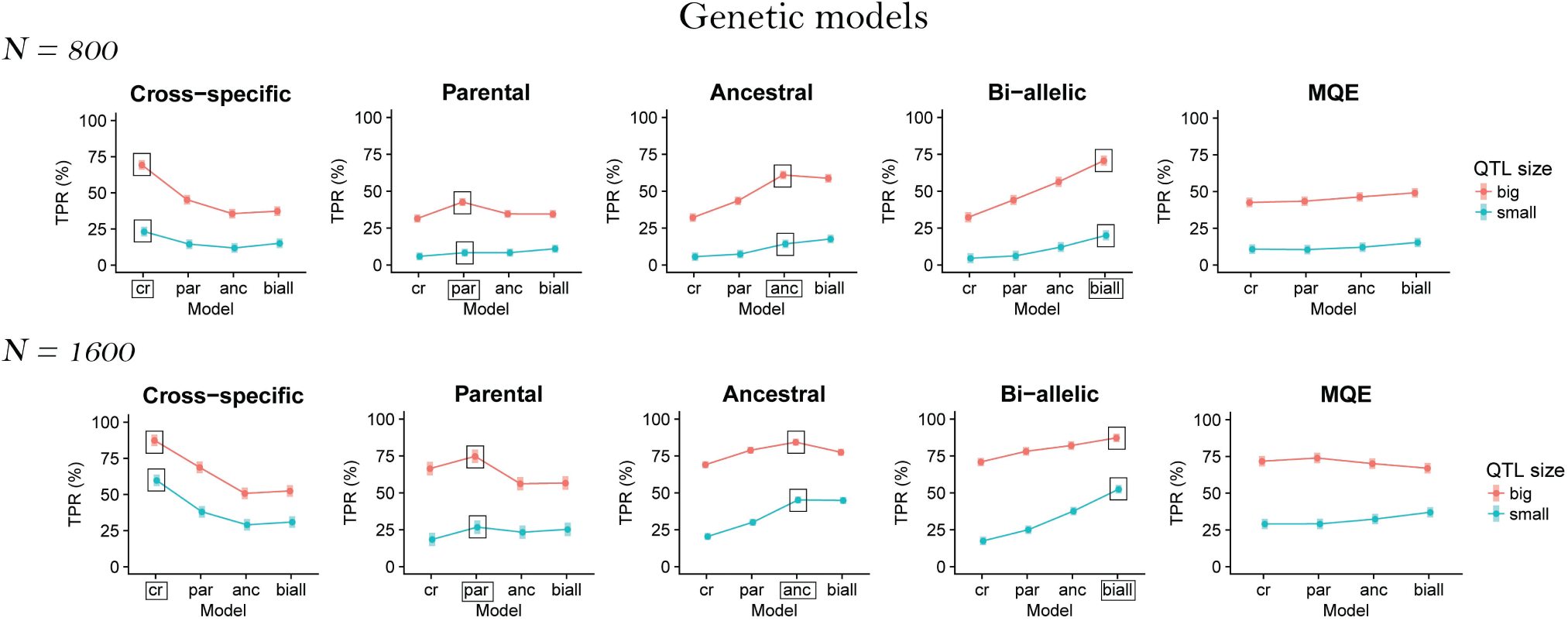
TPR LSM over the QTL detection model (cross-specific, parental, ancestral, and bi-allelic) and the QTL size (small 2% and big 6%) for all QTL genetic models (cross-specific, parental, ancestral, bi-allelic, and MQE) per population size (N = 800 and N = 1600). The framed results represent the QTL detection model corresponding to the simulated QTL genetic model.

## IV. DISCUSSION

The main objective of this article was to evaluate the influence of the QTL allelic diversity on MPP QTL detection. We simulated QTLs with different levels of allelic diversity and evaluated the QTL detection power in different MPP designs (diallel, chessboard, factorial, NAM), derived from five or nine parents. Changing the number of parents produced variation in the covered genetic diversity and the number of individuals per cross. We also varied the total population size, the size of the QTL effect, and the QTL detection model used. We evaluated jointly the effect of all parameters in a statistical model to determine their relative contribution, and we determined CI for the detected QTL positions.

### MPP design

According to the ANOVA (Tables 5 and 6) and the LSM results (Figure 3), the MPP design was mostly important for the cross-specific QTLs. For those QTLs, designs with a reduced number of large crosses like the NAM performed better than designs with many small crosses like the diallel. Obviously, cross-specific QTL effects need large cross sample sizes to be detected. Therefore, to detect QTL effects that are potentially diverse and cross-specific we recommend using designs with few large crosses. Sampling a reduced number of allele genetic background interactions increases the sample size to estimate those effects and also the chance to reject the null hypothesis because we only need to detect a single interaction different from zero. However, the QTL detection and the estimation of the allelic effects should be distinguished. For example, an MPP design with few crosses will be efficient to detect the QTL but it will not allow the correct estimation of all individual QTL allelic effects.

The MPP design was less important for detecting the other types of QTL allelic effects (parental, ancestral, bi-allelic). Contrary to the cross-specific QTL effects, the parental, ancestral and bi-allelic QTL allelic effects are consistently defined across crosses, which gives them an increased sample size. The parental, ancestral and bi-allelic QTL alleles reached more easily the critical sample size for detection, which made them less dependent on a particular MPP design. This result is consistent with the conclusion of [8] who noticed that the form of the design did not influence the detection of parental and bi-allelic QTLs.

### Number of parents

The number of parents used represents a trade-off between the number of sampled alleles and the sample size to detect the QTLs. For a fixed population size, MPP designs using more parents cover a larger genetic diversity but with a reduced number of individuals per cross. Generally, the TPR decreased with a larger set of parents and reduced cross sizes (Figure 4). This result was observed for all types of QTL effects in the N=800 populations and for the QTLs with a small effect (2%) in the N=1600 populations. The use of MPP designs with large crosses is therefore an important factor for MPP QTL detection. This result is consistent with several simulations [10, 11, 16, 6] and empirical cross-validation studies [22].

In MPPs composed of crosses, most of the QTL variance is probably due to the contrast between the parental alleles within the crosses. Moreover, the use of a QTL detection model with a cross-specific intercept accounts for an important part of the between crosses QTL variance [6, 12]. We prefer sampling strategies that increase the sizes of the segregating crosses. In our case, five parents covered probably a sufficient part of the genetic diversity. Therefore resources are better spent on enlarging the sizes of the crosses. This finding is consistent with the conclusions of [8] who showed that QTL detection power reached a plateau after six parents. Therefore, assuming that few parents are enough to sample genetic diversity, MPP designs should rather invest resources to increase the cross sizes than increasing the covered genetic diversity.

Still, in some situations, we did find that using nine parents instead of five increased the TPR. This result was consistent with [8, 16, 3, 12]. In our case, we emphasized that using a larger set of parents was only useful to increase the TPR of QTLs with a low MAF (Figure 5). The parental QTL Q4 and ancestral QTL Q6, which only segregated in a single parent and ancestral group respectively, and the low MAF bi-allelic QTLs were the only cases where using nine parents increased the detection power. We hypothesize that when the QTL MAF was low, using more parents enabled to sample at least one cross where the QTL segregated. However, we noticed that the TPR only increased in the large populations (N = 1600) and for the big QTLs. Therefore, the use of more parents should be combined to an increased total population size and will be conditioned on the QTL effect size. A large population and a big QTL effect increase the chance of detection when only few individuals carry the QTL allele.

The results concerning the number of parents illustrate the interest to consider the question of the resource allocation in MPPs given the QTL allelic diversity. In general, we observed that larger crosses increased the TPR for all types of QTL effects in the N=800 populations. The negative effect of cross size reduction was less important for QTLs with shared effects between crosses like the ancestral QTLs. For those QTL alleles, the reduction of the within cross sample size was compensated by an increased between cross sample size. However, in the larger populations (N=1600), we noticed that increasing the number of parents to cover a larger genetic diversity could be useful to detect rare QTLs with a large effect. According to [23], those QTLs effects represent an important part of the genetic variation and are good candidates for MAS. Therefore, in recurrent MAS breeding programs with population size between 800-1000, we advise to prioritize the enlargement of the cross sizes. However, if the objective is to enrich material with new traits coming from diverse material, it might be worthy to increase the covered genetic diversity and the total population size to detect influential alleles.

### Mapping resolution

The results about the MPP design and the number of parents showed a difference between type of QTL allelic effects. The FDR and dQTL results also reflected this distinction. In Table 3, we noticed that the cross-specific and parental QTLs were detected with larger FDR and dQTL than the ancestral and bi-allelic QTLs. The increased resolution and reduced FDR observed for the ancestral and bi-allelic QTLs could again be due to an increased sample size to detect the consistent QTL allelic effects.

The FDR decreased when the tolerated distance to the simulated QTL increased. For example, around 60% of the QTLs detected as false positive at 10 cM become true positive if we enlarge the tolerated distance to 20 cM. Many false positive could be explained by the extent of the linkage disequilibrium, which is relatively large in *F*_2_ populations. The FDR indicates that, in MPPs composed of *F*_2_ crosses, the CI around a detected QTL should be wide to include the true QTL position. For example, in Table 4, we showed that the 95% empirical CI of the 2% QTLs detected in the N = 800 MQE populations, was equal to 70 cM. For the 6% QTLs the 95% empirical CI was 46 cM. Using a CI of at least 50 cM around the detected QTLs seems to be necessary in MPPs composed of *F*_2_ crosses. This information is a first attempt to increase the reliability in the selected region around a detected QTL in MPPs composed of crosses, which is an important criteria for MAS [5].

### Simulation validity

The low FDR on the chromosome with no simulated QTLs confirms that our QTL detection procedure functioned properly. Extrapolating FDR chr to the whole genome gave us values between 0.9 and 15.3% with an average of 5.9%. These values are comparable to the type I error of 10% of the empirical thresholds that supported our choice [19].

We also compared the TPR we obtained with results from the literature. For example, [24] estimated the QTL detection power of interval mapping method in *F*_2_ populations with 10 QTLs accounting for a total of 30% of the phenotypic variation. This scenario was the closest to our simulation settings. In that case, [24] used a threshold with type I error of 25% and he obtained powers of 57% and 85% for total population sizes of 500 and 1000 individuals, respectively. We compared those values to the TPR obtained with the bi-allelic genetic model scenario detected with the bi-allelic model in the N=800 populations because this scenario was the closest to the one used in [24]. We looked at the TPR with no minimum distance to the simulated QTL obtained in the simple interval mapping scan. We used − *log*10(*p* − *value*) = 3 as significance threshold to account for the larger type I error used by [24]. In that case we obtained a TPR value of 69% comparable to the 57 – 85% interval obtained by [24] for 500 and 1000 individuals, which supports the credibility of our simulation.

### Application to other types of populations and species

The simulations we performed were based on *F*_2_ crosses starting from real sugar beet data. Our main conclusions were similar to the one of authors that used different type of populations. Indeed, [11] who used *F*_2_, backcross (BC), and full-sibs populations and [16] who used double haploid (DH) reached the same conclusion about the importance of using large crosses. Therefore, we can reasonably consider that our conclusions could generalize to other type of populations.

In our simulations, we assumed that all genotypes in a cross came from the same *F*_1_ plant. However, as emphasized by a reviewer, depending on the crop, it might not be possible to generate *F*_2_ crosses containing 450 genotypes from a single *F*_1_ cross. If both parents are inbred, the *F*_1_ will be identical. However, if the parents are partially heterozygous, as it was the case for three of our parents, then *F*_1_ genotypes might differ. The most problematic situation occurs when the same marker is homozygous in one *F*_1_ and heterozygous in another. In that case, if the genotypes are mixed in the same cross, the marker will segregate in parts of the cross and be monomorphic in the rest. Then, the origin of the monomorphic marker will be wrongly assigned to a parent when this origin is actually unknown. So far, the best solution we have found for this problem is to specify the sub-cross structure due to the use of different *F*_1_ plants. This situation happening in many breeding populations must be taken into consideration. The R package we used to perform the QTL detection includes such an option in the data processing.

The generalization of our results to other crops would need further confirmations. The use of simulated data starting from real genotypes is a strategy that has been used in several articles [25, 15, 26]. Few of those studies have tested hypotheses about the trade-off between the number of crosses and the number of individual per cross. Among them, [26] did not find a significant influence of the cross size in simulated MPPs from rapeseed genotypes. The total population size (2000) and the number of simulated QTLS (50) were however different from our simulation settings. In [3], the authors simulated 25-100 QTLs on a real maize NAM population with 5000 individuals. They found that increasing the number of parents (crosses) increased the QTL detection power. Assuming that many of the simulated QTLs had a low MAF, this result would be consistent with our second conclusion: increasing the number of parents is beneficial to detect QTLs with low MAF if the total population size is large enough to guarantee a minimum cross size. Since those results represent too few comparison points, we would need other studies to infer about the generalization of our results to other crop species.

## V. CONCLUSIONS

We tried to determine the most powerful MPP design to detect QTL in MPPs given various levels of QTL allelic diversity. We could show that the trade of between using a large set of parents to cover a larger genetic diversity with smaller crosses and reducing the number of parents to increase the cross-size depends on the type of QTL allelic effects. In most of the cases, sampling a reduced number of parents is enough to cover a sufficient amount of the genetic diversity. Therefore, resources should be used to increase cross sizes rather than increasing the covered allelic diversity. However, when the main goal is the detection of QTLs with rare allele and large phenotypic effects, using a larger set of parents can be beneficial given that the total population size is also increased. Concerning the QTL detection resolution in MPPs composed of *F*_2_ crosses, we noticed that those populations have a low resolution. We advise to use a CI of at least 50 cM around the detected QTL positions to have a reasonable probability that it includes the true QTL position.

