## Supplementary material for "The influence of QTL allelic diversity on QTL detection in multi-parent populations: a simulation study in sugar beet"

February 2, 2020

### S1: Illustration of the simulated QTL effect on the reference diallel

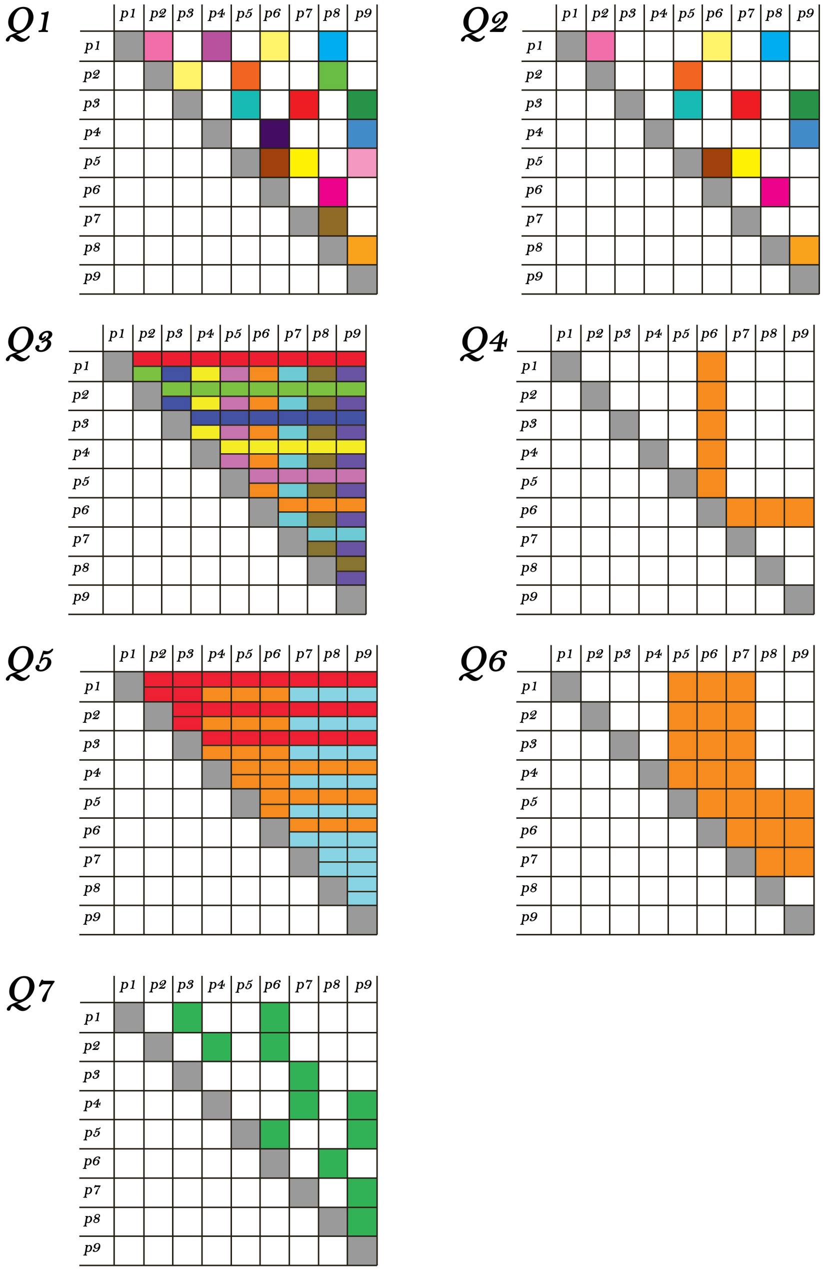


Figure 1: Q1: cross-specific QTL segregating in half of the crosses, Q2: cross-specific QTL segregating in one third of the crosses, Q3: parental QTL with non-zero allelic effect for every parents, Q4: parental QTL with non-zero allelic effect for a single parent, Q5: ancestral QTL with non-zero allelic effect for all ancestors, Q6: ancestral QTL with non-zero allelic effect for a single ancestor, Q7: bi-allelic QTL

#### S2: Phenotypic values simulation procedure

For each realizations, phenotypic values were simulated at the level of the reference diallel population (36 crosses with 450 genotypes). The $i^{th}$ realization included the following steps:

1. The phenotypic variance was expressed in terms of QTL and random error variance $\sigma_p^2 = \sigma_Q^2 + \sigma_e^2$. We assumed a strict additivity of these components.

2. We randomly sampled the 8 QTL positions $(q_1, ..., q_8)$. Each QTL was on a different chromosome. We assumed that the QTL positions were independent and that the global QTL variance $(\sigma_Q^2)$ was the sum of each individual QTL variance contribution $(\sigma_Q^2 = \sum_{i=1}^{n_{QTL}} \sigma_{qi}^2)$. We calculated the individual QTL variance using $\sigma_{qi}^2 = V(\boldsymbol{X_{q_i}\beta_i})$ where $X_{q_i}$ and $\boldsymbol{\beta_i}$ are the incidence matrix and the allelic effect of QTL $i$. The incidence matrix $X_{q_i}$ took different forms according to the type of simulated QTL effect (cross-specific, parental, ancestral, bi-allelic). The form of $\boldsymbol{\beta_i}$ followed the definition of the simulated QTLs (Q1-7). The non-zero elements of $\boldsymbol{\beta_i}$ were sampled from a uniform distribution (1-10) with random sign assignment. We scaled the $\boldsymbol{\beta_i}$ values to make sure that the $\sigma_{qi}^2$ reached the desired phenotypic proportion (2 or 6 %). Finally we calculated the QTL contribution to the phenotype using $\boldsymbol{y_Q} = \boldsymbol{X_Q\beta}$.

3. We determined the error variance contribution $(\sigma_e^2)$ such that $\sigma_e^2 = ((1 - h^2)/h^2) * \sigma_Q^2$. In all cases, $h^2$ was equal to 0.32. Given $\sigma_e^2$ we sample the phenotypic variation due to the error using $\boldsymbol{y_e} \sim N(0, \sigma_e^2)$. The simulated phenotypic values were therefore expressed as $\boldsymbol{y_{sim}} = \boldsymbol{y_Q} + \boldsymbol{y_e}$.

4. In many cases, even if the QTLs were sampled on different chromosomes, there was still an important covariance between the QTL positions. Therefore, we sampled a large number of realizations and we kept only the one where the covariance between the QTL positions was inferior to 1% of the total phenotypic variance.

### S3: Histograms of the distance between simulated and detected QTL distributions

The red, blue, and black dashed lines represent the .90, .95 and .99 distribution quantiles.

## N = 800

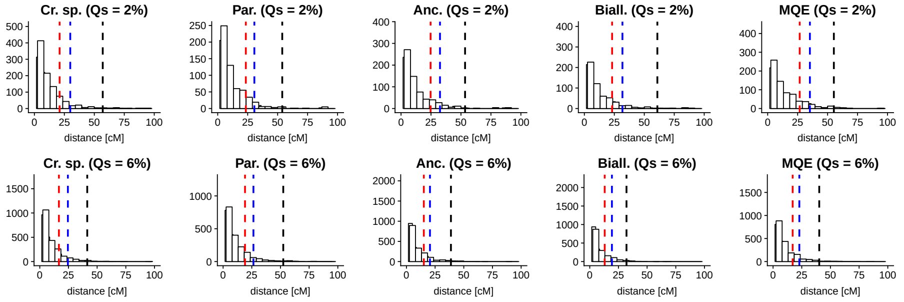


## N = 1600

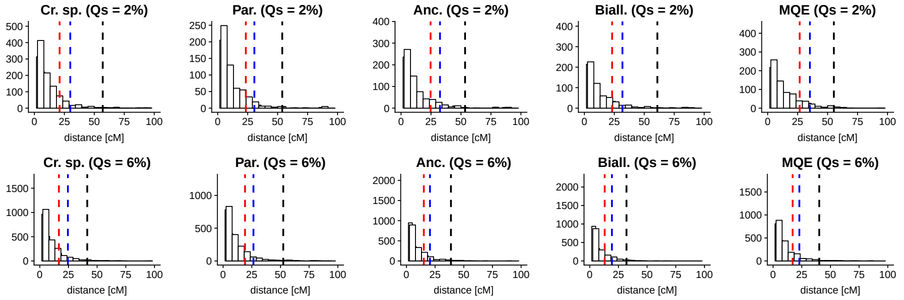
